# Psychophysical validation of calibrated white-noise and tuned-noise auditory masks for transcranial focused ultrasound

**DOI:** 10.64898/2026.07.29.741617

**Authors:** Martin T.W. Scott, Patti N. Limon, Gerald Popelka, Kim Butts Pauly, Anthony M. Norcia, Ryan T Ash

**Affiliations:** University of California, San Francisco; Stanford University

## Abstract

Auditory confounds have proven to be a major hurdle in the elucidation of veridical neuromodulation effects with transcranial ultrasound stimulation (TUS). Auditory noise masks are an essential method to reduce the audibility of TUS and have shown promise in several studies. Here we describe a novel approach for design, calibration, and psychometric validation of auditory noise masks to reduce the perceptibility of TUS. White noise masks and spectrum-tuned masks matched to a TUS protocol that generates highly salient auditory costimulation (487.5 Hz pulse repetition frequency, 10% duty cycle, 68 W/cm2 pulse-peak average intensity, 500 kHz acoustic frequency) were generated, and dB(A) levels were calibrated with an artificial ear. The masker levels needed to reduce TUS detection performance in a two-interval forced choice task were determined with an adaptive QUEST+ staircase in 20 neurotypical participants. Detection performance of this highly salient TUS protocol was driven to near chance performance (<55%) in 17/20 participants with white-noise and 18/20 participants with spectrum-tuned noise. However, high masker levels approaching safety limits were needed to render TUS inaudible for the majority of participants, indicating the need for formal masker calibration for these TUS settings.

Additionally, against expectation the spectrum-tuned masker did not significantly outperform the white-noise masker, suggesting that perceptibility of TUS auditory costimulation does not lawfully follow the sound expected from its pulse envelope and the known spectrum of human hearing.

## Introduction

Transcranial ultrasound stimulation (TUS) is an emerging neuromodulation method with great potential for use in causal manipulation studies in neuroscience and neuropsychiatric therapeutics owing to its high spatial precision and depth penetration. Although ultrasound is defined as sound with frequencies above the range of human hearing, the ultrasound signal is often pulsed at audible frequencies (5 - 500Hz) in search of pulse repetition frequency- (PRF-) dependent effects, rendering the TUS audible (Gavrilov 2014). The perceived pitch and loudness of the audible components are determined by several properties of the TUS protocol: the PRF; the pulse intensity/pressure, the duty cycle, and shape of the envelope (i.e. ramped pulses are less audible than square-wave pulses (Mohammadjavadi et al. 2019, Johnstone et al 2021). Additionally, different depths of stimulation can sometimes be audibly distinguishable when matching the intensity at the focus, because deeper sites require a higher output intensity. This auditory costimulation may lead to confounds. Subjective awareness of TUS violates experimental blinding because participants and experimenters can easily differentiate active from control conditions. Auditory costimulation can bias participants by distracting or attracting attention to some conditions over others, and it can generate neural responses in auditory circuits that are spuriously attributed to the ultrasound stimulation itself (Guo et al. 2018; Sato et al. 2018; Guo et al. 2023). Recently, auditory costimulation was found to explain suppressive effects of TUS on the transcranial magnetic stimulation (TMS) motor evoked potential (Kop et al. 2024). Therefore, successfully elimination of auditory costimulation is a key need.

Previous studies have attempted to reduce the impact of auditory costimulation by adding an external auditory masking signal. Most experimenters attempt to match the spectral content of the mask to the spectral content of the assumed audible TUS pulse envelope (Braun et al. 2020; Eraifej et al. 2026; Johnstone et al. 2021; Butler et al. 2022), or to combine a white noise mask with a PRF matched masker (Kop et al. 2024; Braun et al. 2020). A more recent approach is to use a complex multi-tone sound to camouflage the ultrasound tones (Liang et al. 2023; Strohman et al. 2024; Legon et al. 2024). This is consistent with the principle of critical-band theory, which predicts that masker energy near the sound-to-be-masked’s frequency contributes most strongly to masking, whereas energy outside the ‘critical band’ contributes little additional masking. In other words, by using a spectrally tuned mask, effective masking should be possible with a quieter (and thus more tolerable) sound.

However, this benefit comes with a trade-off: in experiments in which multiple PRFs are used, a different auditory mask is needed for each PRF, making the PRF conditions of the experiment audibly distinct. This may complicate the interpretation of results, as the PRF condition has been effectively un-blinded, and necessitates controls for this un-blinding.

Another potential downside is that any sounds generated by the TUS that are not predicted by the PRF and DC will not be masked by a tuned masker. A simpler approach is to use a white-noise mask. A white noise mask has equal energy at all frequencies and therefore has the potential to adequately mask any combination of ultrasound parameters. Additionally, a white noise mask is technically simple to implement. However, the white noise’s uniform spectrum also implies that most of the mask spectrum is noncontributory to masking the audible TUS components, but still contributes to the perceived loudness of the mask.

Therefore the white noise may need to be at a high level that is unpleasant to the participant or may even exceed safety limits for the inner ear. However, if a white noise mask can perform adequately at acceptable levels, it offers the advantage of being fixed across experiment conditions.

To inform experimental design decisions for transcranial focused ultrasound, the purpose of the present experiment was to compare the masking effectiveness of a white noise mask and a ‘spectrally tuned’ noise masker.

## Methods

### Participants

Twenty neurotypical participants aged 18-55 (mean=33), 11F, 9M, were recruited from the community to participate in the study. All participants completed informed consent under a protocol approved by the Stanford University Institutional Review Board.

### Ultrasound parameters

TUS was administered using a 500KHz 4-channel transducer (NeuroFus CTX-500) with a TPO-203 4-channel drive system. The parameters used were always: PRF = 487.5 Hz, spatial peak pulse average intensity (I_SPPA_) = 68 W/cm^2^, duty cycle = 14%, pulse duration = 0.287 ms, pulse train = 0.960ms, beam-steering depth = 70mm. These parameters were chosen to be a close match to ongoing experiments in our laboratory.

### Auditory maskers

Two different masks were used in this experiment: a white noise mask and a spectrally tuned mask. The masks were synthetically generated with Python software. For both masks, the duration was 4.8s, and the sampling rate was 192 kHz. The white noise mask was constructed in the time domain by taking samples from a Gaussian distribution. The tuned mask started as a white noise sound generated in the same way, but it was then shaped in the frequency domain to more closely match the harmonic structure expected from a rectangular pulse train with a fundamental frequency of 487.5 Hz and a duty cycle of 14% (that is, the spectral structure of the audible TUS components). The filter used to shape the white noise was constructed as the sum of multiple Gaussian frequency bands centred on the integer harmonics of the PRF (487.5 Hz, 975.0 Hz, 1462.5 Hz, and so on). For each PRF harmonic frequency the equivalent rectangular bandwidth (ERB) was calculated, and this value was used to set the full-width half-maximum (FWHM) for the associated Gaussian frequency band. The ERB approximates the integration bandwidth of human hearing at audible frequencies. The ERB was calculated following from Glasberg and Moore (1990), and was used as the FWHM of the Gaussian, giving:

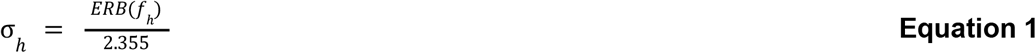

Where *ERB* is equation 3 from Glasberg and Moore (1990) and *f*_*h*_ is the frequency of the harmonic *h*. The scale of the Gaussian distribution was set according to the spectral envelope produced by using a 14% duty cycle, which can be obtained from the cardinal sine (sinc) function:

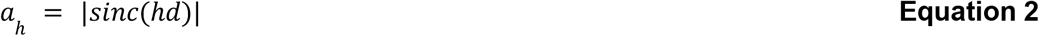

Where *a* _*h*_ is a scaling coefficient for the Gaussian distribution associated with harmonic *h* and duty cycle *d*. For each harmonic, using the associated σ_*h*_ and *a*_*h*_ terms, the Gaussian distribution was evaluated at every frequency *f*_*k*_, where *k* is the spectral bin index, and *f*_*h*_ is again the frequency of harmonic *h*:

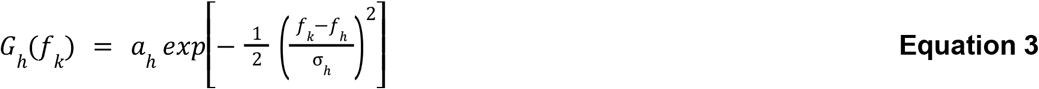

Where the limit of *k* is the spectral bin of the sound’s Nyquist limit. Then, the complete filter was formed by summing these Gaussian distributions across all harmonics:

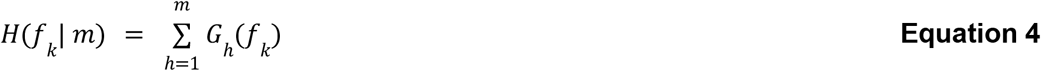

Where the maximum harmonic *m* was determined as the point at which 10 consecutive harmonics had an *a*_*h*_ beneath a cutoff criterion of −40dB relative to the peak amplitude at the

PRF. This prevents the generation of gaussian distributions at frequencies with amplitude so low as to not be audible, reducing mask generation time. The fast Fourier transform (FFT) of the white noise was multiplied by the filter function *H*(*f*) and inverse transformed back into the time domain. The spectra of both masking sounds used is overlaid with the calculated 487.5Hz 14% duty cycle TUS spectrum in **Figure 1**. The white noise masker has constant spectral density across all frequency bins, whilst the spectrally tuned masker has peaks at the frequencies present in the actual TUS sound, particularly at lower frequencies where the ERB is relatively narrow. Note, as the harmonic frequency grows, so does the ERB, such that neighbouring harmonics have overlapping ERBs (this is noticeable starting from approximately 2 kHz in **Figure 1**).

**Figure 1.**
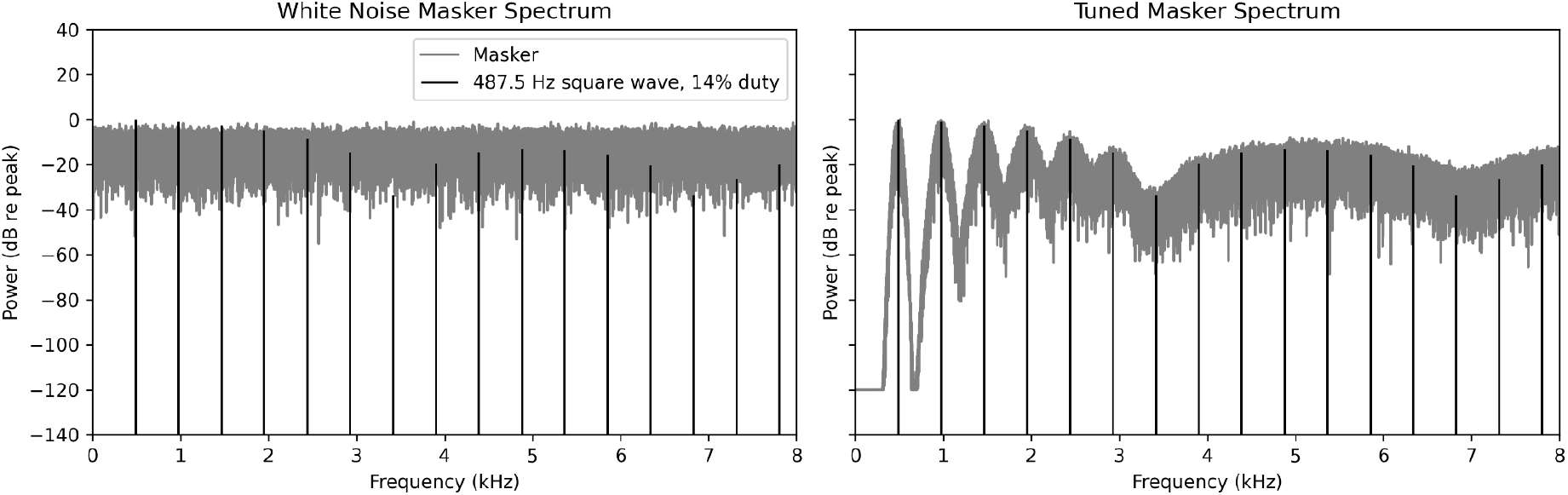
illustration of rationale for matched-spectrum masker, spectra for white noise and matched spectrum masker

### Mask level calibration

Both masker sounds were delivered over a pair of earphones (Koss KSC75, https://koss.com/products/ksc75). These earphones were calibrated using a sound level meter (Larson Davis SoundAdvisor 831c) with ear-simulating earphone coupler (Larson Davis AEC201-A). For each mask sound, 15 loudness levels were used, starting from approximately 63.5 dB(A), to a maximum of 91 dB(A) (steps of approximately 2 dB(A)). We chose a maximum value of 91dB(A) based on sound exposure dose guidelines from The National Institute for Occupational Safety and Health (NIOSH), which allows for 2 hours of continuous exposure at 91 dB(A). We reasoned that 2 hours is more than enough time for most TUS experiments that will involve exposure to an auditory mask (in our laboratory, the actual TUS delivery portion of an experiment is rarely over ~35 minutes). Additionally, we chose to use conventional earphones rather than bone-conduction headphones to avoid the added complexity of calibrating bone-conducted output.

### Procedure

Participants were positioned in a brain stimulation experiment chair (Rogue Research, Inc.), and their hair was prepared with ultrasound gel above the left ear following published best practices (Murphy et al. 2025). We did not use neuronavigation to position the transducer for this experiment, though the placement above the left ear with a 10-degree angled gel pad was an approximate match to the location used in our ongoing focused ultrasound experiments of the visual thalamus (lateral geniculate nucleus). The earphones were placed on the participant’s ears bilaterally. Participants were then instructed on and practiced the task. The temporal sequence of a task trial is illustrated in **Figure 2**. In each trial, the ultrasound pulses could occur in one of two short (960ms) intervals, with each interval separated by 50ms of mask-only (that is, two-interval forced-choice). During each trial, either the white noise mask or the tuned mask was played. The order of masking conditions was pseudo-randomised per-participant. After the second interval, participants had 1.2 seconds to press a button on a keypad (either button 1 or button 2) to report whether the TUS occurred during the first or second interval. Following each trial, the level of the mask was updated with a QUEST+ Bayesian adaptive algorithm (Watson 2017). There was a separate QUEST+ staircase initiated for each masker condition. The QUEST+ algorithm was set to search for the threshold and slope of the psychometric function underlying the 2IFC detection of TUS, which we assumed to be a standard Weibull function. The lower asymptote was fixed to 50% (2IFC guess rate), and the lapse rate to 2%. The possible mask level values and possible threshold values were set to the 15 calibrated loudness levels specified earlier, and the possible slope values were 12 geometrically spaced values between 1 and 38.9. Participants completed 125 trials per auditory mask condition, leading to a total of 350 trials. All participants were subject to 5-10 practice trials during which no mask was played, allowing them to form an understanding of the sounds and sensations associated with TUS exposure. Participants practiced until they correctly responded to at least three consecutive no-mask practice trials. Then, they were subject to 5-10 more practice trials with masking sounds played over headphones. The experiment took approximately 45 minutes.

**Figure 2.**
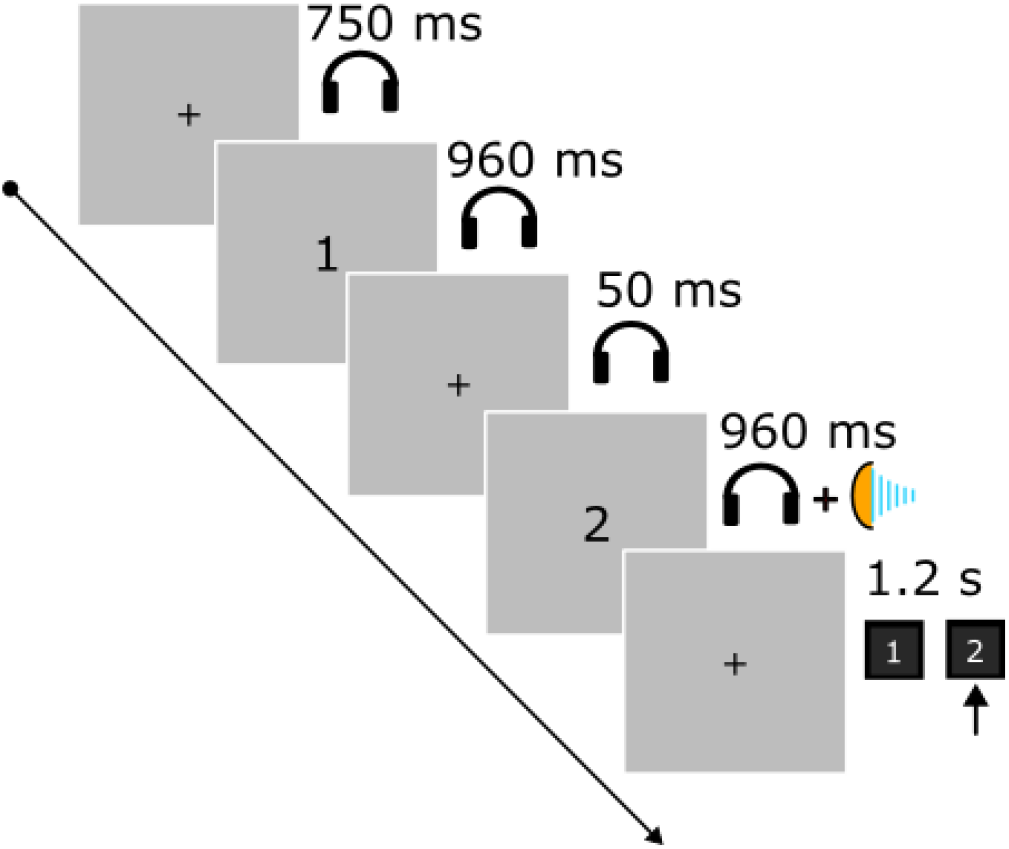
Illustration of a single trial in the experiment. Each trial began with onset of an auditory masker (shown by headphones). Two 960 ms TUS intervals separated by a 50 ms interval were indicated by numbers displayed on the screen. TUS was administered in one of these two intervals (TUS icon shows TUS administered in second interval in this example). TUS parameters were 487.5 Hz PRF, 68 W/cm2 I_SPPA_, 14% duty cycle, 70 mm Z-depth steering, 500 kHz acoustic frequency. Following the second interval, the participant had 1.2 seconds to report the interval in which TUS occurred using a keypad. The level of the mask was determined on each trial with a QUEST+ adaptive staircase (see Methods). The order of mask conditions (white noise or tuned noise) was pseudo-randomized across trials.

### Psychometric function fits and statistics

For each participant and masker condition, a Weibull psychometric function was fitted to the trial-level data (Python toolbox psychofit, https://github.com/cortex-lab/psychofit) using maximum-likelihood optimization. Data fits were based on absolute masker level in dB(A), but the fitting was implemented on a reversed internal axis so that larger internal x-values corresponded to quieter maskers and therefore higher detection performance. Only the threshold and slope parameters were estimated, and the asymptote values were fixed to that used by the QUEST+ procedure. In actual dB(A) units, the fitted threshold range was constrained to 60-100 dB(A), while the slope range was constrained to 0.1-80.0. For each participant, the fitted Weibull model was used to predict the probability of detecting the TUS at the maximum permitted masker loudness (91 dB(A)). TUS detectability at 91 dB(A) was compared between the white and tuned masker using a two-sided exact paired sign test applied to participant-level fitted detection probabilities. The within-subject difference (white noise - tuned noise) was computed for each participant, and the sign test evaluated whether the direction of these paired differences was equally likely to favor either masker condition under the null hypothesis; exact zero differences were treated as ties to be excluded (there were no ties).

## Results

Example QUEST+ adaptive staircase psychometric curves for both masker types used are shown for a representative participant in **Figure 3**. Percentages correct at different mask levels are shown, with the number of trials at the intensities sampled by QUEST+ indicated by the symbol size. The arrows on these plots indicate that this participant would have a 52.5% chance of hearing the TUS for a white noise masker at 91 dB(A) and a 50% chance using the tuned masker. This detectability at 91 dB(A) was calculated for all participants for both maskers and used group-level analysis. The 91 dB(A) level was the maximum masker level employed due to NIOSH safety limits (see Methods).

**Figure 3.**
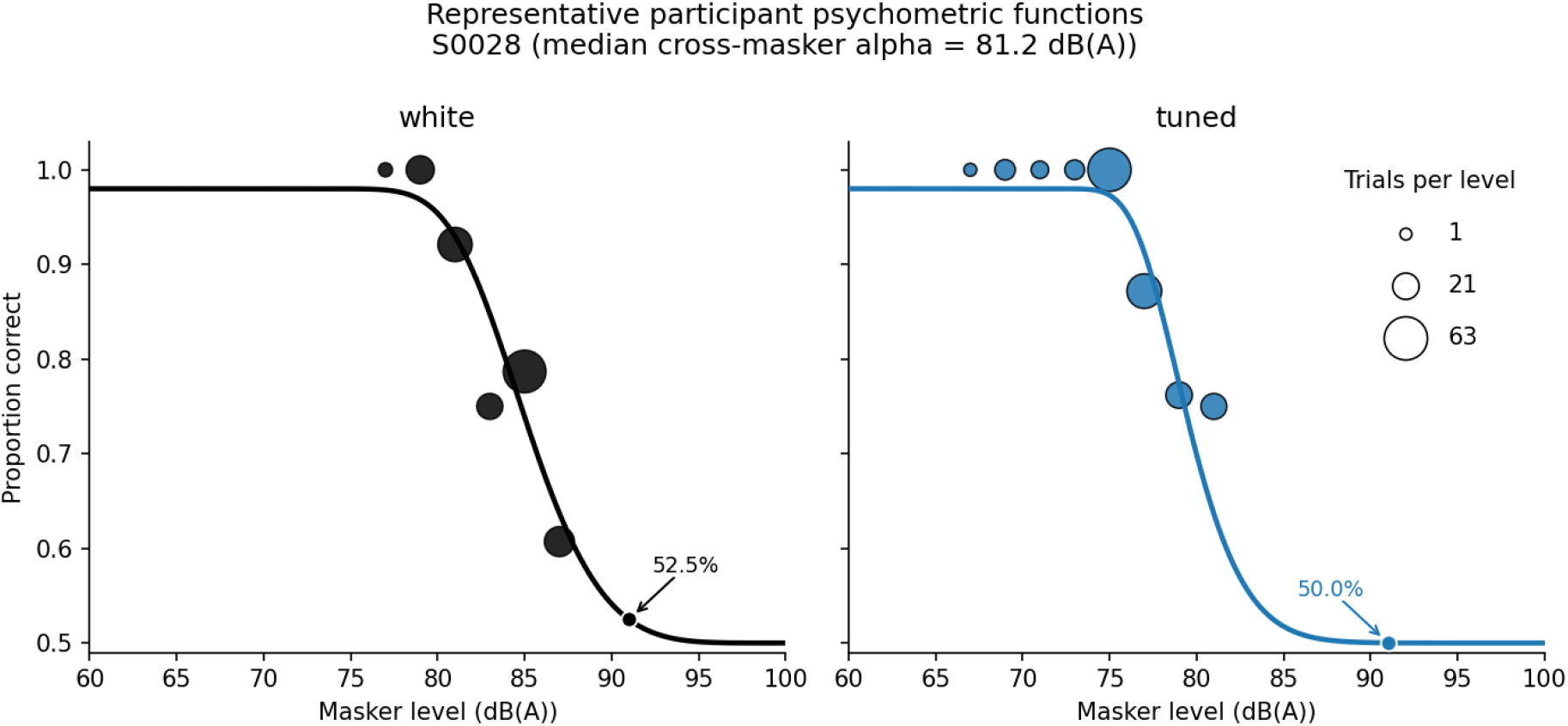
An example Weibull psychometric function fit for a participant with median performance in the sample. The size of the symbols indicates the number of trials at a given masker level. The arrow indicates the intercept between the model fit and a masker level of 91dB(A).

In **Figure 4**, we present the group-level data in two different ways: the probability of TUS detection at 91 dB(A) (the maximum masker level used in the experiment, panel A), and the modelled mean sensitivity at each masker level based on the average across-participant psychometric functions (panel B). Panel A shows that the majority of participants had near-chance detection probabilities (<55%) using a 91 dB(A) masker for both masker types used (17/20 for white noise, 18/20 for tuned noise). There was a trend towards the white noise masker having a slightly higher detectability (mean = 52.4%, SD = 4%) than the tuned noise masker (mean = 51.2%, SD = 2.6%). However, at 91 dB(A), fitted detection probability did not differ reliably between masker conditions by a two-sided exact paired sign test: the detectability of TUS under white noise exceeded tuned noise in 11 participants and tuned noise exceeded white noise in 8 participants (N = 20, no ties), p = 0.648. The mean paired difference (white noise - tuned noise) was 1.19% (in units of detectability).

**Fig. 4.**
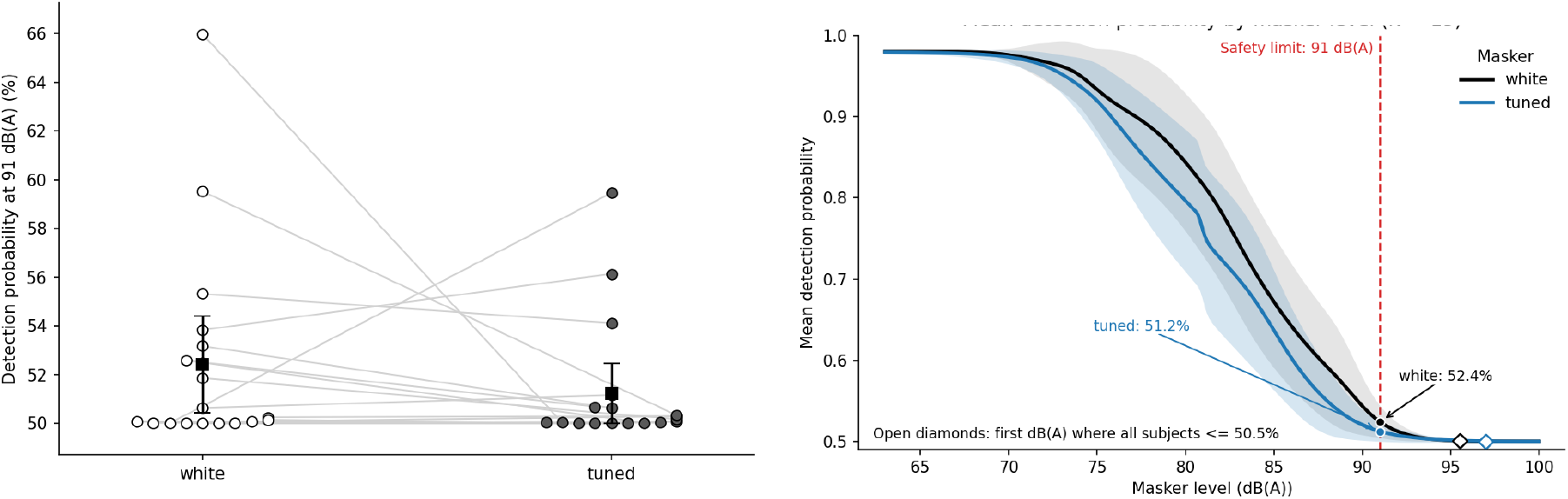
Group data. A. Predicted detection probability at 91 dB(A) for the white noise and tuned noise maskers, including mean (squares) and 95% confidence intervals bracketed lines). Individual participant data points are depicted as circles (open for white noise, filled for tuned noise). **B**. Mean detection probability as a function of masker level in dB(A) constructed from psychometric function fits for white noise (black) and tuned noise (blue).

Shaded regions show 95% CIs. Diamond markers show the predicted loudness required to drive all participants to 50.5% detection probability.

## Discussion

The purpose of this study was to investigate the relative TUS masking effectiveness of white noise and a TUS-matched spectrally tuned noise intended to mask the audible perception of TUS. Across 20 participants, the detectability of a usually clearly audible 487.5Hz PRF focused ultrasound pulse train was dramatically reduced in most cases, with near chance performance (<55% percent correct) achieved in 17/20 participants with white noise and 18/20 participants with tuned noise both masks. The participants with remnant performance were still partially masked, with detection performance of 55-66% with white noise and 56-59% with tuned noise. Surprisingly, there was scant evidence for an improvement in masking performance when using the tuned masker compared to the white noise masker. The only evidence for an improvement with a tuned masker was the presence of a larger effect size for the tuned masker when comparing the mean 51% threshold to our 91 dB(A) limit, but this did not bear out in direct statistical comparisons of the maskers we used. To our knowledge, no other publication has tested TUS masking using calibrated signals. Thus, no other report can accurately state noise levels required for masking in terms of audiometric units with a notion of the safety and maximum acceptable duration of sound exposure.

One reason that may explain why the white and tuned maskers had similar performance is the method used to determine the placement of the tuned-mask Gaussians. As the harmonic frequency increases, the width of the ERB increases too, meaning the Gaussians overlap progressively more, and the tuned-mask spectrum begins to resemble a white-noise masker spectrum. If the audibility of the TUS is most apparent at these higher-harmonics (where the white noise mask and tuned mask are more similar), it therefore may be unsurprising that the tuned mask offers less of an advantage as most of the ‘tuning’ exists between 0 and 3.5 kHz where the ERBs overlap far less. It is likely that the advantage of a tuned masker manifests more at lower PRFs (5Hz) where the majority of the audible content occurs at frequencies with narrower ERBs, and thus a greater potential for ‘tuning’. It is also possible that filtering the mask spectrum on the basis of ERBs is contributing to these findings. An alternative approach would be to restrict the widths to be much narrower, such that more of the masker energy is delivered closer to the harmonic present in the sound being masked.

Future researchers can explore whether tuning ERB width reveals a clear advantage of using a tuned masker over a white noise masker. Another potential reason for the lack of difference between the two maskers is that the auditory perception generated by the transducer is not lawfully predicted by the envelope wave shape. It’s possible that the electronics of the transducer generate other sounds unrelated to the pulse envelope, eliminating the benefit of the tuned masker. Overall, our findings indicate that the trade-off for using a tuned masker may not be worth the added complexity in generation, calibration, and loss of conditional blinding when using multiple PRFs or DCs.

Several other studies have tested spectrally tuned auditory masks and demonstrated efficacy. Braun et al. (2020) demonstrated that a square wave masker matched to the TUS PRF and duty cycle (PRF 1 kHz, duty cycle 50%, other parameters: acoustic frequency 500 kHz, pulse duration 300 ms, max pressure in brain < 0.6 MPa) can, in the majority of participants, drive participants’ ability to detect active from control TUS to chance. The mask also eliminated the TUS-induced auditory event-related potential from scalp electrodes.

However, the authors did not report how they calibrated the level of the mask or provide an estimate of the mask level, nor did they report the free-water transducer output in I_SPPA_ or MPa, making direct comparison to our paradigm (in terms of masker and TUS audibility) difficult. The same absence of reporting limitations apply in a subsequent study by the same authors (Butler et al. 2022), where the efficacy of the same 1 KHz square wave masker was reported in a new cohort of 16 participants (PRF 1 kHz, DC 50%, pulse duration 300 ms, max pressure in brain < 0.6 MPa, acoustic frequency 500 kHz), and shown to drive TUS detectability (d’) to near zero, on average. Johnstone et al. (2021) used a similar masking approach with a square-wave masker matched to the PRF and duty cycle (PRF 250 Hz, DC 50%, acoustic frequency 270 kHz, free-water I_SPPA_ 16 W/cm^2^, pulse duration 300 ms, with and without envelope ramping). Calibrating the masker level to “the limit of tolerability”, these authors were able to effectively mask the TUS detectability in half (3/6) of participants. It is important to note that these authors used a lower acoustic base frequency of 270 kHz, which produces more pronounced somatosensory costimulation compared to 500 kHz (Kop et al. 2025), which may explain their difficulty in masking effectively.

Generally, in experiments that have assessed masking rigorously, there is an absence of masker level measurement and reporting, making explorations of differences in masking efficacy difficult. Nevertheless, masking efficacy in our study appeared better or equivalent to that in previous reports. In Braun et al., despite group-level masking success, 3/18 participants (17%) remained clear outliers who could detect masked TUS, with d′ values >3. In Butler et al., group-level d′ was close to zero, but individual values were variable; visual inspection of the reported d′ values suggests that approximately 5/16 participants (25%) had residual positive discriminability above d′ ≈ 0.5 in the their control site condition with a maximum around 1.29. In Johnstone et al., PRF-matched headphone masking reduced but did not eliminate audibility, with approximately 3/6 participants still able to detect TUS above 60% in the masked condition. Compare this to our results in which no participants achieved a high ability to detect TUS: 2 of 20 participants >55% correct, 1 of 20 participants (5%) with d’ > 0.5 (max d’ = 0.58).

Notably, one masker type that has not yet been rigorously assessed with actual ultrasound exposure is complex multitone auditory mondrian mask (Liang et al. 2023). Publications that employed this masking method have instead opted to use a questionnaire approach at the conclusion of the experiment, and set the masker level to “a level where participants could not hear normal conversations” (Legon et al. 2024; Strohman et al. 2024). While this approach allows for a gross assessment of TUS detectability, it is not sufficient for cross-study comparisons of masking efficacy, as there is potential for expectancy effects confounding participant’s self-report, and the spectrum of speech is highly variable and broadband.

In the present experiment, we treated 91 dB(A) as the level limit, which allows for 2 hours of continuous exposure. According to NIOSH, the time-dosage maximum halves with every 3 dB(A) added to the sound level. Our results show that the majority of participants were driven to chance-level detection with a 91 dB(A) masker (for both a white and tuned mask), but that some still had above-chance detection. To capture all these participants, our psychometric modelling predicted that a masker level of approximately 96 dB(A) would be required. This sound level is only safe for 37 minutes and would be very unpleasant. Thus, unless masking efficiency is considerably increased, these subjects will not be fully masked under the TUS parameters we have used. However, many of the participants above the criterion level are still likely being acceptably masked for experimental purposes, where 18 out of 20 participants have a <55% chance of detection. Notably, there was also a great deal of variance in the masker level required to produce a near chance-level performance. In this dataset, the standard deviation was approximately 10 dB(A) (a 10 dB(A) addition in volume is perceived as roughly twice as loud). This suggests that calibrating masker level based on the individual experimenter’s experience may be a poor representation of masking effectiveness across the population of participants. This recapitulates the need to rigorously verify masker effectiveness under different TUS experimental conditions. In particular, when the only control condition is a sham no-TUS condition (i.e. there is no active control site), participant unblinding can easily happen. Even in the case of an active neural TUS site, differences in audibility between the beam-steered positions, or differences in the physical position of the transducer, could cause cryptic unblinding of participants, which can lead to small but statistically significant differences in behavior or neural activity across conditions that can be spuriously attributed to neuromodulation by ultrasound (Guo et al. 2018; Kop et al. 2024). We note here that the PRF we chose (487.5 Hz) generates highly salient auditory perceptions, which in pilot studies was the most difficult to mask. We therefore chose this PRF for our experiment as a worst-case scenario. Lower PRFs that are commonly used in TUS studies, e.g. 5-10 Hz, will be able to be masked at significantly lower masker levels, which could be made even lower by ramping the onset and offset of the TUS envelope to decrease the level of higher harmonics. Ramping was not viable at the PRF tested here, because the pulses are so brief (0.28 ms).

Our study has important limitations that will need to be expanded upon in the future. First, we only tested spectrum-matching for a single TUS protocol, with a single PRF and TUS intensity. Therefore our masker characteristics do not directly apply to most existing TUS studies. TUS experiments with lower PRFs and lower intensities, and/or pulse ramping, will be able to use lower masker levels that are more comfortable to participants. Another limitation is that the tuned mask was likely not optimized. A different tuning spectrum would possibly have masked the TUS sound more efficiently. This is a task for future work which will improve the safety and comfort of auditory masks. A third limitation is that this auditory masking will not reduce the perceptibility of TUS tactile perception. Although at 500 kHz acoustic frequencies, tactile perception is minimal, at lower acoustic frequencies tactile perception becomes clearly perceptible (Kop et al. 2025). Masking this perception would require more complex interventions beyond those described here.

In conclusion, here we describe a method for calibration of a white noise or spectrum-tuned auditory mask and rigorous confirmation of mask efficacy in TUS experiments. Our results show that even highly salient TUS auditory perception can be safely masked with a white-noise auditory mask, and that ERB-based spectral shaping of the auditory mask does not significantly improve masking compared to white noise. The methods are readily implementable in TUS research labs. A white noise stimulus is easily generated with existing software tools.

We hope that this and related work supports adequate auditory masking and improves the validity and reproducibility of TUS experiments in the coming years. This work provides a foundation for rigorous confirmation of masker effectiveness in future TUS studies.

## References

Braun, Verena, Joseph Blackmore, Robin O. Cleveland, and Christopher R. Butler. 2020. “Transcranial Ultrasound Stimulation in Humans Is Associated with an Auditory Confound That Can Be Effectively Masked.” Brain Stimulation 13 (6): 1527–1534.

Butler, Christopher R., Edward Rhodes, Joseph Blackmore, et al. 2022. “Transcranial Ultrasound Stimulation to Human Middle Temporal Complex Improves Visual Motion Detection and Modulates Electrophysiological Responses.” Brain Stimulation 15 (5): 1236–1245.

Eraifej, John, Jake Toth, Jeremy Hanemaaijer, et al. 2026. “Suppression of Pathological Oscillations with Transcranial Focused Ultrasound in Parkinson’s Disease.” Nature Communications 17 (1). 10.1038/s41467-026-70714-7.

Gavrilov, Leonid R. Use of focused ultrasound for stimulation of various neural structures. New York, NY, USA: Nova Biomedical, 2014.

Glasberg, B. R., and B. C. Moore. 1990. “Derivation of Auditory Filter Shapes from Notched-Noise Data.” Hearing Research 47 (1-2): 103–138.

Guo, Hongsun, Mark Hamilton 2nd, Sarah J. Offutt, et al. 2018. “Ultrasound Produces Extensive Brain Activation via a Cochlear Pathway.” Neuron 98 (5): 1020–1030.e4.

Guo, Hongsun, Hossein Salahshoor, D. Wu, et al. 2023. “Effects of Focused Ultrasound in a ‘Clean’ Mouse Model of Ultrasonic Neuromodulation.” iScience 26 (12): 108372.

Johnstone, Ainslie, Tulika Nandi, Eleanor Martin, Sven Bestmann, Charlotte Stagg, and Bradley Treeby. 2021. “A Range of Pulses Commonly Used for Human Transcranial Ultrasound Stimulation Are Clearly Audible.” Brain Stimulation 14 (5): 1353–1355.

Kop, Benjamin R., Linda de Jong, Butts Pauly Kim, Hanneke E. M. den Ouden, and Lennart Verhagen. 2025. “Parameter Optimisation for Mitigating Somatosensory Confounds during Transcranial Ultrasonic Stimulation.” Brain Stimulation 18 (4): 1224–1236.

Kop, Benjamin R., Yazan Shamli Oghli, Talyta C. Grippe, et al. 2024. “Auditory Confounds Can Drive Online Effects of Transcranial Ultrasonic Stimulation in Humans.” eLife 12 (RP88762): RP88762.

Legon, Wynn, Andrew Strohman, Alexander In, and Brighton Payne. 2024. “Noninvasive Neuromodulation of Subregions of the Human Insula Differentially Affect Pain Processing and Heart-Rate Variability: A within-Subjects Pseudo-Randomized Trial.” Pain 165 (7): 1625–1641.

Liang, William, Hongsun Guo, David R. Mittelstein, Mikhail G. Shapiro, Shinsuke Shimojo, and Mohammad Shehata. 2023. “Auditory Mondrian Masks the Airborne-Auditory Artifact of Focused Ultrasound Stimulation in Humans.” Brain Stimulation 16 (2): 604–606.

Mohammadjavadi, Morteza, Patrick Peiyong Ye, Anping Xia, Julian Brown, Gerald Popelka, and Kim Butts Pauly. 2019. “Elimination of Peripheral Auditory Pathway Activation Does Not Affect Motor Responses from Ultrasound Neuromodulation.” Brain Stimulation 12 (4): 901–910.

Murphy, Keith R., Tulika Nandi, Benjamin Kop, et al. 2025. “A Practical Guide to Transcranial Ultrasonic Stimulation from the IFCN-Endorsed ITRUSST Consortium.” Clinical Neurophysiology: Official Journal of the International Federation of Clinical Neurophysiology 171 (March): 192–226.

Sato, Tomokazu, Mikhail G. Shapiro, and Doris Y. Tsao. 2018. “Ultrasonic Neuromodulation Causes Widespread Cortical Activation via an Indirect Auditory Mechanism.” Neuron 98 (5): 1031–1041.e5.

Strohman, Andrew, Gabriel Isaac, Brighton Payne, Charles Verdonk, Sahib S. Khalsa, and Wynn Legon. 2024. “Low-Intensity Focused Ultrasound to the Insula Differentially Modulates the Heartbeat-Evoked Potential: A Proof-of-Concept Study.” Clinical Neurophysiology: Official Journal of the International Federation of Clinical Neurophysiology 167 (November): 267–281.

Watson, Andrew B. 2017. “QUEST+: A General Multidimensional Bayesian Adaptive Psychometric Method.” Journal of Vision 17 (3): 10.

